# From Exception to Rule: Widespread Production of Methylated Rhamnolipids in the *Burkholderia cepacia* Complex

**DOI:** 10.64898/2026.10.01.756062

**Authors:** Maude Dagenais Roy, Marie-Christine Groleau, Xavier Perron, Eric Déziel

**Affiliations:** Centre Armand-Frappier Santé Biotechnologie, Institut National de la Recherche Scientifique (INRS), Laval, Québec, Canada

**Keywords:** Surfactant, Biosurfactant, Glycolipids, *rhlM*, Methyltransferase, Structural diversity, Bioluminescence screening

## Abstract

Although *Burkholderia cenocepacia* exhibits biosurfactant-dependent phenotypes and possesses an *rhl* operon, the chemical identity of its surface-active product has long remained unresolved. The *rhl* operon of *B. cenocepacia* harbors an additional gene, *rhlM*, predicted to encode a methyltransferase, and recently linked to methylated rhamnolipid synthesis in a few *Burkholderiaceae*. Here, we hypothesized that this biosurfactant remained cryptic due to low expression under standard laboratory conditions. Using a chromosomal P*rhl*-*luxCDABE* fusion reporter, we screened diverse nutritional conditions and identified a defined medium containing galactose and urea that strongly induced *rhl* transcription. Liquid chromatography–tandem mass spectrometry (LC-MS/MS) analyses of *B. cenocepacia* K56-2 extracts identified the biosurfactants as methylated dirhamnolipid congeners, predominantly L-rhamnosyl-L-rhamnosyl-3-hydroxytetradecanoyl-3-hydroxytetradecanoate methyl ester (RR-C_14_-C_14_Me). Extending our analysis across the *Burkholderia cepacia* complex (Bcc) revealed that methylated rhamnolipid production is widespread and restricted to *rhl*-positive species. Comparative genomic and protein structural analyses further demonstrated high conservation of both the *rhlM*-containing operon architecture and the RhlM protein across the Bcc. Together, our findings identify the long-standing cryptic biosurfactant of *B. cenocepacia* as methylated rhamnolipids and establish rhamnolipid methylation as a conserved hallmark across the Bcc, shifting the paradigm of rhamnolipid structural diversity from an isolated exception to a widespread rule.

**IMPORTANCE:** The chemical identity of the long-standing cryptic biosurfactants produced by the *Burkholderia cepacia* complex (Bcc) has remained unresolved due to low expression under standard laboratory conditions. Using a bioluminescent reporter screening approach, we identified the responsible operon and defined specific nutritional parameters that promote biosurfactant production, showing that *Burkholderia cenocepacia* synthesizes methylated rhamnolipids. Our findings establish rhamnolipid methylation as a conserved and functionally significant feature among *rhl*-positive Bcc members, mediated by the conserved RhlM methyltransferase. This work reframes the paradigm of rhamnolipid structural diversity, proposing that methylated congeners may constitute the ancestral glycolipid form. Beyond expanding our understanding of secondary metabolism in opportunistic pathogens, this study highlights RhlM as a promising biocatalytic tool for the engineering of custom non-ionic biosurfactants.

## INTRODUCTION

The *Burkholderia cepacia* complex (Bcc) comprises closely related *Burkholderia* species found in diverse environmental niches and recognized as opportunistic pathogens (1). Their ability to thrive in such diverse environments relies in part on the production of a wide array of secondary metabolites, a trait shared by many members of *Burkholderia sensu lato* (2). Among these, several Bcc species have been reported to release, or exhibit phenotypes associated with the production of, surface-active metabolites, often called biosurfactants (3–11). These versatile amphiphilic molecules are synthesized by various microorganisms and reduce surface tension (12). However, the chemical nature of such compounds apparently produced by Bcc bacteria has historically remained cryptic.

Among the different classes of biosurfactants, rhamnolipids represent potential candidates for these unidentified surface-active compounds, as genes involved in their biosynthesis are widely distributed among Bcc members (13). Rhamnolipids are glycolipids that comprise a hydrophilic head made of rhamnose sugar moieties and a hydrophobic tail composed of fatty acids (14). Their biosynthesis typically involves three key enzymes, RhlA, RhlB, and RhlC, encoded by the *rhlA*, *rhlB*, and *rhlC* genes, respectively, which are commonly organized within one (e.g., in *Burkholderia thailandensis*) or two (e.g., in *Pseudomonas aeruginosa*) *rhl* gene clusters (7, 13, 15). First, RhlA synthesizes the lipidic, hydrophobic part of the molecule by linking two hydroxylated fatty acids to form 3-(3-hydroxyalkanoyloxy)alkanoic acids (HAA) (16, 17). Then, the rhamnosyltransferase RhlB adds a first rhamnose moiety to the HAA backbone, forming a monorhamnolipid (18). Finally, a second rhamnosyltransferase, RhlC, can add a second rhamnose to the monorhamnolipid to form a dirhamnolipid (19).

Rhamnolipids display considerable structural diversity, with more than 60 different congeners described (20). These congeners can differ in the number of rhamnose moieties, the length and degree of unsaturation of their fatty acid chains, or the presence of additional functional groups (20). Moreover, a single bacterial species usually produces several congeners, resulting in characteristic rhamnolipid profiles that vary among producers (20, 21).

Beyond the core biosynthesis genes, some *rhl* biosynthetic clusters contain additional genes that may further contribute to rhamnolipid structural diversity (13). Notably, an additional gene named *rhlM*, located between *rhlA* and *rhlB*, is particularly prevalent within the *Burkholderiaceae* family, with a high occurrence in the *Burkholderia sensu lato* genus, especially among members of the Bcc (13, 22). RhlM appears to act as a methyltransferase and, accordingly, methylated rhamnolipids were recently discovered in cultures of the Bcc *Burkholderia lata* 383 (23) as well as *Paraburkholderia kirstenboschensis* F3 (22). In the latter study, methylation of the rhamnolipid was confirmed to be performed through RhlM (13, 22). These findings revealed an additional level of structural diversity among rhamnolipids and suggested that methylated congeners may have been overlooked in other bacteria carrying *rhlM* (13).

Several lines of evidence suggest that the well-studied Bcc species *Burkholderia cenocepacia* produces a rhamnolipid-like biosurfactant. Indeed, we previously reported that knocking out *rhlA* in *B. cenocepacia* K56-2 abolishes biosurfactant-associated phenotypes, including oil emulsion capacity and swarming motility (11). Surfactant activity was reported in the environmental isolate *B. cenocepacia* BSP3, where the molecule responsible was classified as a glucolipid but was not structurally characterized (10). More recently, colorimetric assays have suggested rhamnolipid production by *B. cenocepacia* K56-2 (24). However, these assays do not allow for structural resolution and are susceptible to carbohydrate interference (25). Thus, despite genetic, phenotypic, and biochemical evidence supporting rhamnolipid production, the chemical identity of the biosurfactant produced by *B. cenocepacia* has remained unresolved.

Here, we hypothesized that tailored nutritional conditions would upregulate the *rhl* operon of *B. cenocepacia* K56-2, allowing for the characterization of its cryptic biosurfactant product. Using a transcriptional reporter system, we identified culture conditions that strongly induced the expression of the *rhl* biosynthetic operon as a proxy to obtain better production of the unknown biosurfactant. This allowed us to reveal that *B. cenocepacia* K56-2 indeed synthesizes rhamnolipids and that these molecules are methylated, providing a likely explanation for why they had not previously been identified. Finally, we demonstrate that rhamnolipid production is a functionally conserved hallmark widespread among species belonging to the Bcc.

## MATERIALS AND METHODS

### Bacterial strains and culture conditions

Bacterial strains and plasmids used in this study are listed in **Tables 1 and 2**, respectively. Unless otherwise specified, bacteria were routinely cultivated from -80°C glycerol stocks in Tryptic Soy Broth (TSB) (Difco) in a TC-7 roller drum (New Brunswick Scientific) overnight at 37°C at 150 rpm, or on Lysogeny broth agar (LBA) plates.

**Table 1.** Bacterial strains used in this study.

| STRAIN | RELEVANT CHARACTERISTICS | REFERENCE |
| --- | --- | --- |
| ***Escherichia coli*** |  |  |
| SM10 | Conjugative donor strain, Km<sup>R</sup> | (26) |
| Trans1-T1 | Chemically competent strain used for plasmid cloning | TransGen Biotech |
| ***Burkholderia cenocepacia*** |  |  |
| K56-2 | Clinical isolate; WT | (27) |
| K56-2::pKnock-tc-*rhIA* | *rhIA* insertion mutant; pKnock-Tc integrated into *rhIA*; Tc<sup>R</sup> | (11) |
| K56-2::*rhIA-lux* | K56-2 bearing a chromosomal *rhIA-luxCDABE* fusion reporter | This study |
| LMG19240 | Environmental isolate; WT | (28) |
| HI4437 | Environmental isolate; WT | *Burkholderia cepacia* Research Laboratory and Repository, University of Michigan, USA |
| HI2876 | Environmental isolate; WT | *Burkholderia cepacia* Research Laboratory and Repository, University of Michigan, USA |
| ***Burkholderia ambifaria*** |  |  |
| LMG19467 | Clinical isolate; WT | (29) |
| LMG19182 (AMMD) | Environmental isolate; WT | (30) |
| ***Burkholderia anthina*** |  |  |
| VC16083 | Clinical isolate; WT | Canadian *Burkholderia cepacia* complex Research and Referral Repository, University of British Columbia, Canada |
| HI3538 | Environmental isolate; WT | (31) |
| ***Burkholderia arboris*** |  |  |
| VC8833 | Clinical isolate; WT | Canadian *Burkholderia cepacia* complex Research and Referral Repository, University of British Columbia, Canada |
| ES0263 | Environmental isolate; WT | (32) |
| ***Burkholderia cepacia*** |  |  |
| BTS13 | Clinical isolate; WT | (33) |
| LMG1222 | Environmental isolate; WT | (34) |
| ***Burkholderia contaminans*** |  |  |
| FFH2055 | Clinical isolate; WT | (35) |
| HI4402 | Environmental isolate; WT | *Burkholderia cepacia* Research Laboratory and Repository, University of Michigan, USA |
| ***Burkholderia diffusa*** |  |  |
| VC14008 | Clinical isolate; WT | Canadian *Burkholderia cepacia* complex Research and Referral Repository, University of British Columbia, Canada |
| HI2617 | Environmental isolate; WT | *Burkholderia cepacia* Research Laboratory and Repository, University of Michigan, USA |
| ***Burkholderia dolosa*** |  |  |
| CEP0021 | Clinical isolate; WT | (36) |
| LMG21443 | Environmental isolate; WT | (37) |
| ***Burkholderia lata*** |  |  |
| VC6377 | Clinical isolate; WT | Canadian *Burkholderia cepacia* complex Research and Referral Repository, University of British Columbia, Canada |
| 383 | Environmental isolate; WT | (38) |
| ***Burkholderia metallica*** |  |  |
| VC15467 | Clinical isolate; WT | Canadian *Burkholderia cepacia* complex Research and Referral Repository, University of British Columbia, Canada |
| HI3647 | Environmental isolate; WT | *Burkholderia cepacia* Research Laboratory and Repository, University of Michigan, USA |
| ***Burkholderia multivorans*** |  |  |
| VC17546 | Clinical isolate; WT | Canadian *Burkholderia cepacia* complex Research and Referral Repository, University of British Columbia, Canada |
| LMG17588 | Environmental isolate; WT | (39) |
| ***Burkholderia pyrrocinia*** |  |  |
| LMG21824 | Clinical isolate; WT | (37) |
| CH-67 | Environmental isolate; WT | (40) |
| ***Burkholderia seminalis*** |  |  |
| HI2490 | Environmental isolate; WT | *Burkholderia cepacia* Research Laboratory and Repository, University of Michigan, USA |
| ***Burkholderia stabilis*** |  |  |
| LMG18138 | Clinical isolate; WT | (41) |
| HI2462 | Environmental isolate; WT | *Burkholderia cepacia* Research Laboratory and Repository, University of Michigan, USA |
| ***Burkholderia stagnalis*** |  |  |
| MSMB1956WGS | Environmental isolate; WT | Menzies School of Health Research and Northern Arizona University, USA |
| ***Burkholderia territorii*** |  |  |
| MSMB1502WGS | Clinical isolate; WT | Menzies School of Health Research and Northern Arizona University, USA |
| MSMB301WGS | Environmental isolate; WT | Menzies School of Health Research and Northern Arizona University, USA |
| ***Burkholderia ubonensis*** |  |  |
| LMG24263 | Clinical isolate; WT | (32) |
| LMG20358 | Environmental isolate; WT | (42) |
| ***Burkholderia vietnamiensis*** |  |  |
| VC11253 | Clinical isolate; WT | Canadian *Burkholderia cepacia* complex Research and Referral Repository, University of British Columbia, Canada |
| LMG22486 | Environmental isolate; WT | (43) |

**Table 2.** Plasmids used in this study.

| PLASMID | RELEVANT CHARACTERISTICS | REFERENCE |
| --- | --- | --- |
| mini-CTX-*lux* | Chromosomal integration vector carrying *luxCDABE*; Tc<sup>R</sup> | (44) |
| pXP1 | mini-CTX-*lux* derivative carrying the *B. cenocepacia* K56-2 *rhl* operon promoter upstream of *luxCDABE*; Tc<sup>R</sup> | This study |
| pFTP1 | Plasmid carrying a trimethoprim resistance cassette; Tp<sup>R</sup> | (45) |
| pBBR5::FLP | Broad-host-range plasmid expressing FLP recombinase; Gen<sup>R</sup>, Tc<sup>R</sup> | (46) |
| pXP2 | pBBR5::FLP derivative in which the Tc<sup>R</sup> cassette was replaced with Tp<sup>R</sup>; Gen<sup>R</sup>, Tp<sup>R</sup> | This study |

### Construction of the *B. cenocepacia* K56-2 P*rhl*-*lux* reporter strain

Primers used for the construction of the reporter strain are listed in **Table 3**. The promoter region upstream of *rhlA* (locus tag BURCENK562V_RS14995) was amplified from *B. cenocepacia* K56-2 genomic DNA using primers XPRhlGibF and XPRhlGibR. Amplification was performed using Q5® High-Fidelity DNA Polymerase (New England Biolabs, Ipswich, MA, USA). The mini-CTX-*lux* vector was linearized using BamHI and KpnI FastDigest^TM^ restriction enzymes (Thermo Fisher Scientific, Waltham, MA, USA). PCR and digestion products were separated by agarose gel electrophoresis and purified using the FavorPrep^TM^ GEL/PCR Purification Kit (Favorgen Biotech Corp., Pingtung, Taiwan). The purified DNA fragments were assembled using the pEASY®-Uni Seamless Cloning and Assembly Kit (TransGen Biotech Co., Ltd., Beijing, China). All procedures were performed according to the manufacturer’s protocol. Transformants were selected on medium containing 15 μg/mL tetracycline and further screened for luminescence production. The resulting pXP1 plasmid was introduced into *E. coli* SM10.

**Table 3.**
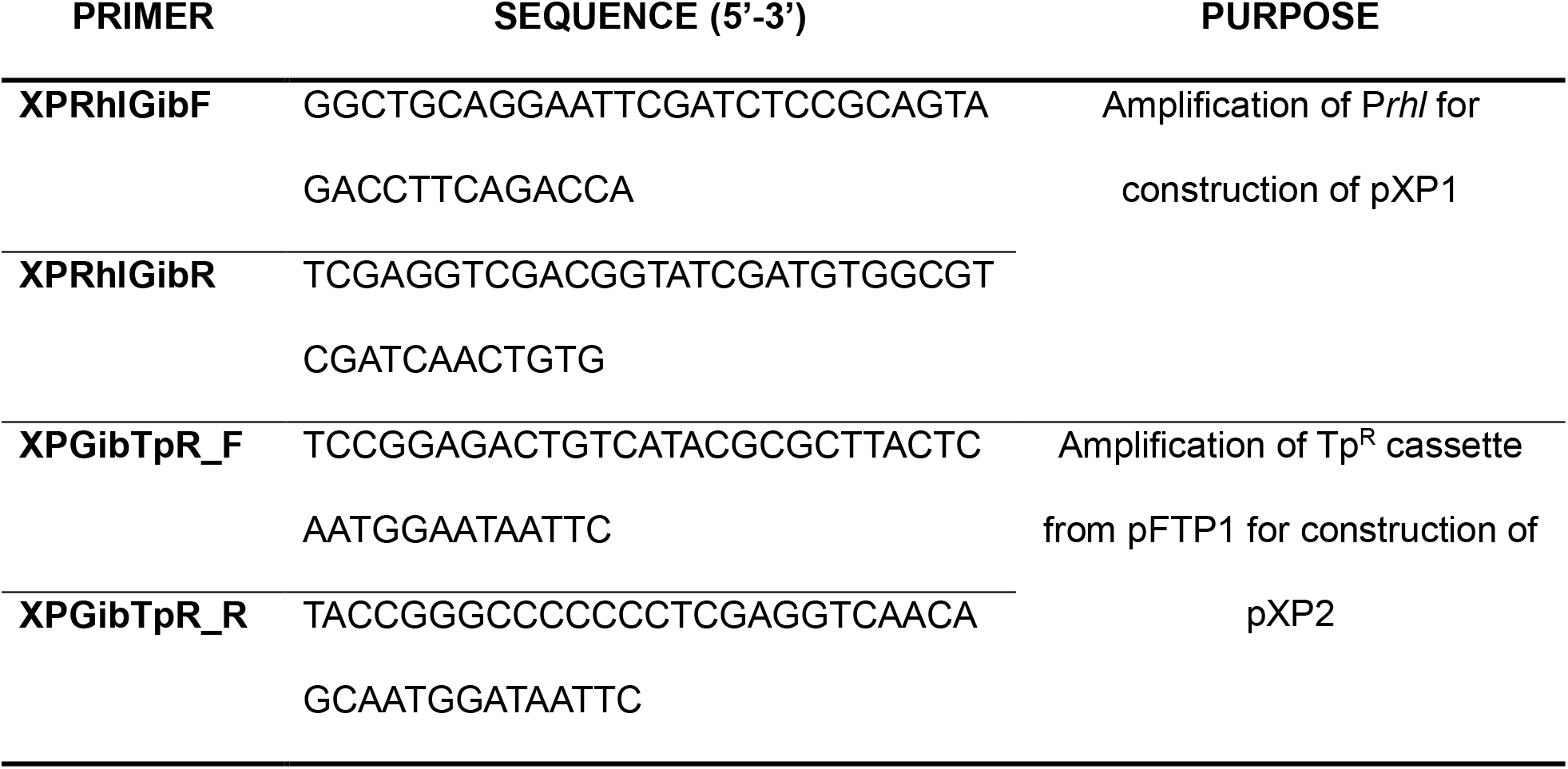
Oligonucleotide primers used in this study.

The pXP1 plasmid was subsequently transferred into *B. cenocepacia* K56-2 by bi-parental conjugation. Aliquots of precultures from each strain were mixed in microcentrifuge tubes at a ratio of 1:1 and plated on TSB plates at 37°C for 24 hours. The cells were recovered from the plate, resuspended in TSB, and 100 μL was plated on selective agar LBA containing 50 μg/mL tetracycline and 15 μg/mL polymyxin B as a counterselective agent against the donor *E. coli* SM10. Plates were incubated at 37°C for 48 hours. Insertions were confirmed using luminescence production.

To remove the FRT-flanked tetracycline resistance cassette following chromosomal integration, the tetracycline resistance gene of pBBR5::FLP was replaced with a trimethoprim resistance cassette amplified from pFTP1 using primers XPGibTpR_F and XPGibTpR_R, generating pXP2. The plasmid was constructed using the seamless cloning procedures as described above. The plasmid was subsequently transferred into *B. cenocepacia* K56-2::P*rhl*::*lux* by conjugation using the procedure described above. Transconjugants were selected on medium containing 200 μg/mL trimethoprim and 15 μg/mL polymyxin B at 30°C for 24 to 48 hours. Colonies were screened for tetracycline sensitivity to confirm FLP-mediated excision of the tetracycline resistance cassette. To eliminate the FLP-encoding plasmid, tetracycline-sensitive colonies were streaked on low-salt LBA supplemented with 10% (w/v) sucrose and incubated at 37°C for 48 h.

### Expression of *rhlA-lux* reporter

To monitor the transcription of the *rhl* operon in different conditions, a *B. cenocepacia* K56-2 reporter strain harboring a chromosomal Pr*hlA*-*lux* fusion was used. For the primary screening, overnight cultures grown in TSB were harvested by centrifugation (10 000 x *g* for 1 minute), washed twice with sterile phosphate-buffered saline (PBS), and resuspended to an OD_600_ of 0.05 in the selected medium. A volume of 200 μL was distributed in 96-well white, clear-bottom microplates. A total of 20 different media formulations (**Table 4**) were tested. Base media included Nutrient Broth (NB), TSB, M8 minimal medium (M9 without NH₄Cl (47); 53 mM Na₂HPO₄, 22 mM KH₂PO₄, 8.6 mM NaCl, 0.3 mM CaCl₂, 1.81 mM MgSO₄, 43 µM FeSO₄, 11 µM MnSO₄, 10 µM ZnSO₄, 3.6 µM CoSO₄), and Basal Salt Medium ((BSM); 18.5 mM K₂HPO₄, 7.25 mM NaH₂PO₄, 0.52 mM nitrilotriacetic acid, 0.05% w/v Casamino acids, and trace metals (810 µM MgSO₄, 43 µM FeSO₄, 11 µM MnSO₄, 10 µM ZnSO₄, 3.6 µM CoSO₄)) and were supplemented with various carbon sources and nitrogen sources at a final concentration of 55.5 mM and 93.5 mM, respectively.

**Table 4.** Culture medium combinations evaluated for *rhl* operon expression in ***B. cenocepacia* K56-2.**

| MEDIUM ID | BASE MEDIUM | CARBON SOURCE | NITROGEN SOURCE |
| --- | --- | --- | --- |
| 1 | NB | Dextrose | Peptone/Beef extract |
| 2 | TSB | Dextrose | Tryptone/Soy peptone |
| 3 | M8 | Dextrose | Ammonium chloride |
| 4 | M8 | Leucine | Proline |
| 5 | M8 | Galactose | Urea |
| 6 | M8 | Sodium pyruvate | Urea |
| 7 | M8 | Lysine | Proline |
| 8 | M8 | Dextrose | Urea |
| 9 | M8 | Fructose | Urea |
| 10 | M8 | Sucrose | Urea |
| 11 | BSM | Dextrose | Ammonium chloride |
| 12 | BSM | Leucine | Proline |
| 13 | BSM | Galactose | Urea |
| 14 | BSM | Sodium pyruvate | Urea |
| 15 | BSM | Lysine | Proline |
| 16 | BSM | Dextrose | Urea |
| 17 | BSM | - | Yeast extract |
| 18 | BSM | Dextrose | Yeast extract |
| 19 | BSM | Leucine | Proline/Yeast extract |
| 20 | BSM | Galactose | Urea/Yeast extract |

Microplates were incubated at 37°C in a Cytation 3 multi-mode plate reader (BioTek Instruments, Winooski, VT, USA). Bacterial growth (OD_600_) and bioluminescence (relative light units, RLU) were simultaneously monitored at 1h intervals over 48 hours, and the plates were agitated for one minute before each measurement. Selected culture conditions identified in the primary screening were subsequently validated in 5-mL cultures in 18*150 mm borosilicate tubes (maintaining a culture-to-tube volume ratio of approximately 1:5 for adequate aeration) incubated in the roller drum. Growth (OD_600_) was monitored using a NanoDrop One Spectrophotometer (Thermo Fisher Scientific, Waltham, MA, USA).

### Oil spraying assay

Biosurfactant production under the most promising culture conditions identified during the screening was further assessed using an atomized oil assay as described by Burch *et al*. (48) and Martinez *et al*. (49). Selected media were solidified with 1,5% (w/v) agar and inoculated with three 5 μ L drops of a *B. cenocepacia* K56-2 suspension adjusted to an OD600 of 3.0. The *B. cenocepacia* K56-2 *rhlA*-mutant was included as a negative control for biosurfactant production. Plates were incubated at 37°C for 48 to 72 hours. Following incubation, the agar surface was sprayed with a fine mist of mineral oil containing 0,5% (w/v) Sudan Red dye. Halos and colony morphologies were visually analyzed and imaged using a binocular microscope (Olympus Life Sciences, Tokyo, Japan), and halo diameters (mm) were measured.

### Quantification of rhamnolipid production in liquid cultures

Bacteria were grown in borosilicate tubes in modified M8 medium supplemented with 55.5 mM galactose and 93.5 mM urea as carbon and nitrogen sources, unless otherwise specified. Cultures were inoculated from overnight cultures at an OD_600_ of 0.05 and were incubated at 30°C for 24 hours in the roller drum at 150 rpm.

High-performance liquid chromatography (HPLC)-grade commercial reagents and solvents were utilized for all extractions without additional purification steps. Whole cultures were extracted thrice with equal volumes of ethyl acetate (Sigma-Aldrich, St-Louis, MO, USA) (1:1, V/V). Before extraction, 2 ppm of the internal standard 5,6,7,8-tetradeutero-4-hydroxy-2-heptylquinoline (HHQ-d4) was added to each culture. The organic phases were pooled, evaporated to dryness under a gentle stream of nitrogen gas, and resuspended in methanol (Sigma-Aldrich, St-Louis, MO, USA) to achieve a 50-fold concentration.

All chromatographic and mass spectrometric analyses were performed using MS-grade solvents and ultra-pure water obtained from a Milli-Q purification system (Merck Millipore, Burlington, MA, USA). The LC-MS analyses were performed with a Quattro Premier XE tandem quadrupole mass spectrometer (Waters Corp., Mildford, MA, USA) equipped with a Z-spray interface using electrospray ionization in the negative mode (ESI-MS/MS). The instrument was interfaced to a Waters 2795 HPLC system (Waters Corp., Mildford, MA, USA) equipped with a Kinetex C_8_ reversed-phase LC column (100 x 4.6 mm, 2.6 μm particle size; Phenomenex, Torrance, CA, USA). LC-MS analyses in scan mode were performed as previously described in Gauthier *et al*. (23).

### Genomic and structural analysis of RhlM homologs

The nucleotide sequence of *rhlM* from *B. cenocepacia* K56-2 (locus tag BURCENK562V_RS14990) was used as a query to identify *rhlM* homologs in Bcc strains for which genome sequences were available using BLASTn (50). The presence and genomic organization of *rhlA*, *rhlM, rhlB*, and *rhlC* were subsequently examined in *rhlM*-positive genomes. The predicted amino acid sequence of RhlM was retrieved and aligned using ESPript 3.0 via its integrated Clustal algorithm (51–53).

Amino acid substitutions relative to the *B. cenocepacia* K56-2 RhlM sequence were identified from the multiple sequence alignment and mapped onto the AlphaFold-predicted structure of *B. cenocepacia* K56-2 RhlM, retrieved from the *Burkholderia* Genome Database (www.burkholderia.com) (54, 55) using PyMOL 2.5.4 (Schrödinger, LLC, New York, NY, USA).

Substitutions were qualitatively classified as conservative or non-conservative based on differences in the physicochemical properties of the corresponding amino acids, including charge, polarity and side-chain characteristics.

### Data processing and statistical analyses

Growth and reporter activity parameters were calculated from OD_600_ and luminescence measurements. The maximum specific growth rate (*μ*_max_) was determined from the exponential phase of the growth curve, and the corresponding doubling time (*G*) was calculated from *μ*_max_. For OD_600_, RLU, and relative luminescence (RLU/OD_600_), total average productivity (P_tot_) was calculated by dividing the final value by the total cultivation time, whereas average productivity at the maximum value (P_max_) was calculated by dividing the maximum value by the time at which it was reached. All experiments were performed in three independent biological replicates. Data are presented as the mean ± standard deviation (SD). Statistical analyses were performed using GraphPad Prism version 11.1.0 (GraphPad Software, Boston, MA, USA). Homogeneity of variances was evaluated using Bartlett’s test. When variances were homogeneous, differences among groups were assessed using ordinary one-way analysis of variance (ANOVA), followed by Tukey’s multiple comparisons test. When variances were unequal, Welch’s ANOVA was used, followed by Dunnett’s T3 multiple comparisons test. Differences were considered statistically significant at *p* < 0,05, with significance levels designated as * (*p* < 0,05), ** (*p* < 0,01), *** (*p* < 0,001), and **** (*p* < 0,0001).

## RESULTS AND DISCUSSION

### Optimization of culture conditions for the expression of the *rhl* biosynthetic genes in Burkholderia cenocepacia K56-2

Although biosurfactant production by *B. cenocepacia* has long been suspected based on several indirect lines of evidence (4, 10, 11, 24), the surface-active molecule produced has yet to be identified. We initially investigated whether *B. cenocepacia* K56-2 produced any of the 60-and- over rhamnolipid congeners previously described in rhamnolipid-producing bacteria, including other *Burkholderia* species (20). However, none of these congeners or any variation in LC-MS peak profiles were detected in K56-2 culture extracts (data not shown) when extracts were compared to those from cultures of the *rhlA*-mutant. We hypothesized that the absence of detection was due to extremely low biosurfactant production levels, below the detection limit of our analytical methods. We thus decided to optimize culture conditions to increase concentrations of biosurfactant produced in cultures and facilitate detection and identification. However, since the molecule remained unidentified, direct quantification was not possible to assess the efficiency of our optimization. To circumvent this constraint, we used a transcriptional reporter strain harboring the promoter region of the *rhl* operon of *B. cenocepacia* K56-2 fused to the *luxCDABE* operon. Luminescence thus served as an indicator of *rhl* operon expression and, presumably, of the unknown biosurfactant production. Using this reporter system, we measured the luminescence to identify culture conditions that promoted higher transcription of the *rhl* operon.

The expression of the *rhl* promoter was screened in 20 different culture medium formulations, encompassing variations in base mineral salts, carbon sources, and nitrogen sources (**Fig. 1A**). NB supplemented with dextrose, which we previously used as the baseline medium for biosurfactant production in *B. cenocepacia* K56-2 (11), served as a reference, along with TSB, also supplemented with dextrose.

**Figure 1.**
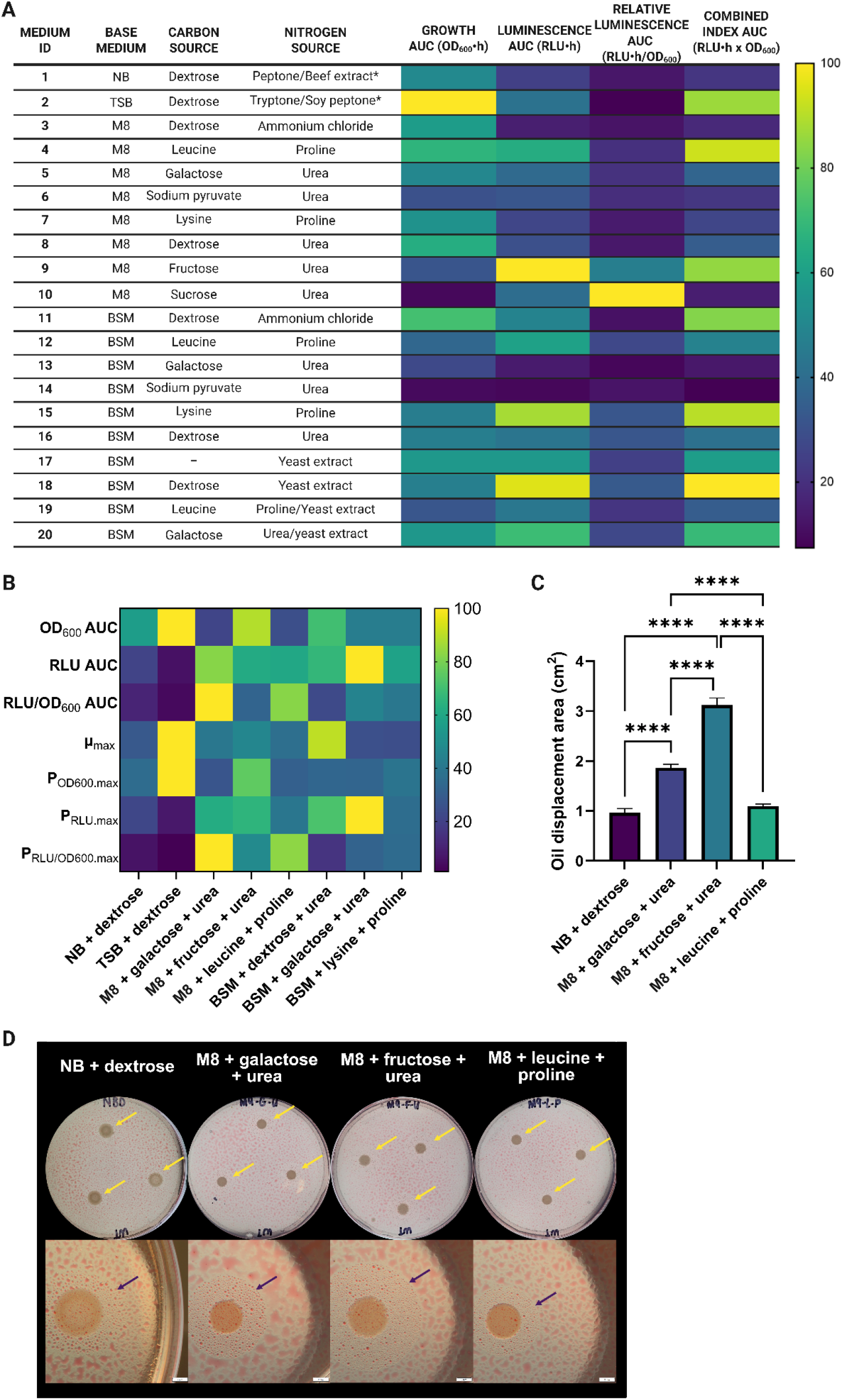
Screening and optimization of culture conditions for *rhl* operon expression and biosurfactant production in *B. cenocepacia* K56-2. (A) Screening of 20 distinct culture media formulations varying in basal salt matrix, carbon sources, and nitrogen sources using a chromosomal *rhlA-lux* transcriptional reporter strain. Growth (OD600), bioluminescence (RLU), and relative P*rhl* activity (RLU/OD600) were monitored over time in 48-hour microplate assays, and the corresponding areas under the curve (AUCs) were calculated. A combined index was calculated as the product of RLU and OD600 to identify conditions where reporter activity and growth compensate for each other. For visualization in the heatmap, AUC values for each parameter were independently normalized to the highest mean value observed across all conditions, which was set to 100%. (B) Characterization of eight selected culture conditions in 5 mL cultures. Growth and P*rhl* activity were monitored over time, and kinetic parameters, including maximum specific growth rate (*μ*max), maximum OD600 productivity, maximum RLU productivity, maximum relative luminescence productivity (RLU/OD600), and AUCs for OD600, RLU, and RLU/OD600, were determined. For visualization in the heatmap, each parameter was independently normalized to the highest mean value observed across the eight conditions, which was set to 100%. (C) Oil displacement activity of four selected culture conditions determined using the atomized oil assay. Oil displacement area was calculated by subtracting the colony area from the total halo area. Data are presented as mean ± standard deviation. Statistical significance was assessed using one-way ANOVA followed by Tukey’s multiple comparisons test. Only selected comparisons are shown (*p* < 0,0001 = ****). (D) Representative images of the atomized oil assay shown in (C), including white-plate images and microscopic visualization of the oil displacement zone. Arrows indicate the boundaries of the displacement halo.

While rich media (**Fig. 1A**; TSB/Medium 2) supported good bacterial growth, only low activity from the *rhl* promoter was detected, which may partly explain why previous identification attempts under these conditions were unsuccessful. The primary screening revealed three critical insights into the nutritional regulation of the *rhl* operon in *B. cenocepacia*.

First, the core mineral base (M8 vs BSM) did not influence the activity of the reporter, demonstrating that defined minimal media, rather than a specific mineral composition, promotes *rhl* operon expression. Moreover, the use of defined minimal media allowed us to better control the culture conditions and evaluate the effect of components on *rhl* operon expression.

Second, within these minimal settings, several sugar-containing formulations led to higher activity of the reporter (**Fig. 1A**). This observation makes sense from a metabolic standpoint, as rhamnolipid biosynthesis relies on precursors derived from central carbon metabolism including dTDP-L-rhamnose (18, 19, 56–58) and hydroxy fatty acids (17, 59, 60). In particular, galactose directly feed into the metabolic pathways that produce glucose-1-phosphate via the Leloir pathway (61), as mapped in *B. cenocepacia* J2315 (KEGG MODULE: M00632; Pathway: bcj00052) (62). Glucose-1-phosphate then serves as the initial substrate for dTDP-L-rhamnose biosynthesis (57, 63). Although these pathways have been historically characterized in *P. aeruginosa*, homologs of the *rmlBDAC* genes responsible for dTDP-L-rhamnose synthesis are widely distributed throughout the Bcc (13, 64), and their enzymatic capability to produce this sugar nucleotide has been biochemically validated in *B. thailandensis* (65).

Furthermore, it is well established that the choice of carbon source plays a critical role in regulating secondary metabolism through carbon catabolite repression (66). Importantly, carbon catabolite repression has also been demonstrated in *B. cenocepacia* K56-2, where glucose and succinate repress the expression of the phenylacetic acid degradation pathway (67). However, carbon catabolite control remains poorly characterized in *Burkholderia* species, where key metabolic signatures have been observed despite the apparent absence of classical homologous genes (68–71). In other rhamnolipid-producing bacteria like *P. aeruginosa*, preferred carbon sources often repress the expression of the *rhl* genes through regulatory systems involving the catabolite repression control protein Crc (72). Thus, differences in carbon catabolite regulation could potentially contribute to the differential *rhl* expression observed among carbon sources in our study.

Moreover, *B. cenocepacia* is a metabolically versatile bacterium commonly found in environmental niches such as the plant rhizosphere (1, 73, 74), Interestingly, several of the nutrients associated with strong *rhl* expression in our study, including simple sugars such as fructose and galactose, as well as amino acids, have been detected in root exudates of plants associated with Bcc species (Fig. 1A) (75–80). The nutrient-dependent regulation of *rhl* expression observed in our study may therefore be relevant to the environmental lifestyle of *B. cenocepacia*, where nutrient availability and composition can vary considerably. In this context, increased biosurfactant production could provide ecological advantages, particularly by facilitating surface-associated motility and interactions with plant roots (12, 81). However, whether the response to specific carbon and nitrogen sources reflects their metabolic utilization or an adaptation to environmental nutrient conditions remains to be determined.

Third, the nature of the nitrogen source strongly influenced reporter activity. Specifically, the presence of urea in the medium generally triggered the most robust and early upregulation of the reporter system, particularly when paired with sugars like galactose, dextrose, fructose, or sucrose (**Fig. 1A**; Media 5, 8, 9, 10, 13, and 16). Interestingly, high reporter expression did not correlate with enhanced biomass accumulation; rather, media containing urea and sugars exhibited moderate to low growth overall compared to those containing ammonium chloride or amino acids (**Fig. 1A**; Media 3, 4, 11). The strong preference for urea as a nitrogen source for *rhl* operon expression could be attributed to both kinetic regulation and pH stability. Indeed, other nitrogen sources, such as ammonium chloride, provide a highly assimilable form of nitrogen that often triggers nitrogen repression, favoring rapid biomass accumulation while silencing secondary metabolism (82, 83). In contrast, urea utilization could bypass global nitrogen repression, creating alternative metabolic dynamics that favor the activation of secondary metabolism at the expense of rapid growth (84–88). Moreover, while the consumption of ammonium ions typically leads to severe medium acidification, urea hydrolysis generates bicarbonate, maintaining a stable neutral pH that preserves enzymatic activity during growth (85, 89, 90).

From the different media evaluated, we selected eight conditions to scale up the experiment, including the two control media, NB and TSB, both supplemented with dextrose (Media 1 and 2), and six of the best minimal media for activation of reporter activity (Media 4, 5, 9, 12, 13, and 16; **Fig. 1A**). These conditions were selected based on their overall performance across growth, cumulative luminescence, and relative P*rhl* activity, combined index for growth and luminescence, as well as the corresponding kinetic profiles (**Supplemental figure S1**). Notably, examining these kinetics and indices allowed us to distinguish true induction from mathematical artifacts, such as in the M8 sucrose-urea medium (**Fig. 1A**; Media 10), where an exceptionally high relative luminescence was driven by slow growth (**Fig. S1C, S1F and S1I**). Three conditions were selected for each minimal medium, M8 and BSM, including different carbon and nitrogen source combinations.

Because the initial screening was performed in 96-well plates, we hypothesized that oxygenation could have been limiting. We therefore tested the selected media in a larger system using 5-mL cultures incubated in a roller drum to increase oxygenation, which is typically crucial for biosurfactant production (91, 92). Our objective here was to select the best-performing medium under our standard laboratory conditions.

Further characterization of the selected media revealed substantial differences in growth and reporter activity (**Fig. 1B**). TSB supplemented with dextrose supported the fastest growth (*μ*_max_ = 0,787 h^-1^; **Table S2** and the highest overall growth but showed low luminescence and relative P*rhl*-*lux* activity. Conversely, defined media supplemented with galactose and urea strongly promoted reporter activity while supporting lower growth. BSM supplemented with galactose and urea generated the highest overall luminescence, whereas M8 supplemented with galactose and urea showed the highest reporter activity relative to cell density (**Fig. 1B**). This indicated that the increased luminescence observed under this condition was not simply attributable to increased bacterial growth, but rather to increased reporter activity per cell.

To determine whether increased P*rhl*-*lux* activity was associated with actual increased biosurfactant production, the eight selected media were also evaluated using the oil spraying assay on agar plates (49). Several defined media produced larger oil displacement zones than our initial reference medium NB supplemented with dextrose (**Fig. 1C-D**). Importantly, this augmented surfactant production was not an artifact of increased bacterial growth or colony expansion, both confounding factors (93). Indeed, while NB supplemented with dextrose yielded the largest colonies (∼9.5 mm), it produced smaller displacement halos than media containing urea, where smaller colony diameters (∼6.2–7.7 mm) consistently produced significantly wider halos (**Fig. 1C-D**, **Fig. S2**). This oil displacement activity was strictly dependent on biosurfactant production, as no halos were observed around the colonies of the *rhlA*-deficient mutant under any of the tested conditions (**Fig. S3**). M8 medium supplemented with fructose and urea showed the largest halo among all conditions, while M8 supplemented with galactose and urea also showed a marked increase compared with NB supplemented with dextrose. Considering these results together with the reporter assays, M8 supplemented with galactose and urea was selected as the optimized condition for subsequent biosurfactant production experiments.

### Identification of the long-standing cryptic biosurfactants in *Burkholderia cenocepacia* K56-2 as rhamnolipid methyl esters

Using these optimized culture conditions, where the expression of the *rhl* operon was maximized, cultures were prepared with the aim of identifying the biosurfactant produced by *B. cenocepacia* K56-2. Culture extracts were then analyzed by LC-MS. An initial analysis in full scan mode was performed to compare the profile of metabolites extracted from wild-type strain cultures vs its *rhlA*-deficient mutant. Because RhlA is strictly required for the very first step of rhamnolipid biosynthesis (16), and the mutant indeed produces no biosurfactant (**Fig. S3**), comparing these two strains allowed us to easily pinpoint the m/z signals that were absent in cultures of the mutant only (**Fig. 2A**). Using this approach, we identified four different peaks dependent on the rhamnolipid biosynthetic machinery, as they were absent in the *rhlA* mutant profile (**Fig. S4**). The predominant congener at *m*/*z* 821.6 matches the profile for L-rhamnosyl-L-rhamnosyl-3-hydroxytetradecanoyl-3-hydroxytetradecanoate methyl ester (RR-C_14_-C_14_Me) we recently identified in cultures *B. lata* (**Fig. 2A**) (23).

**Figure 2.**
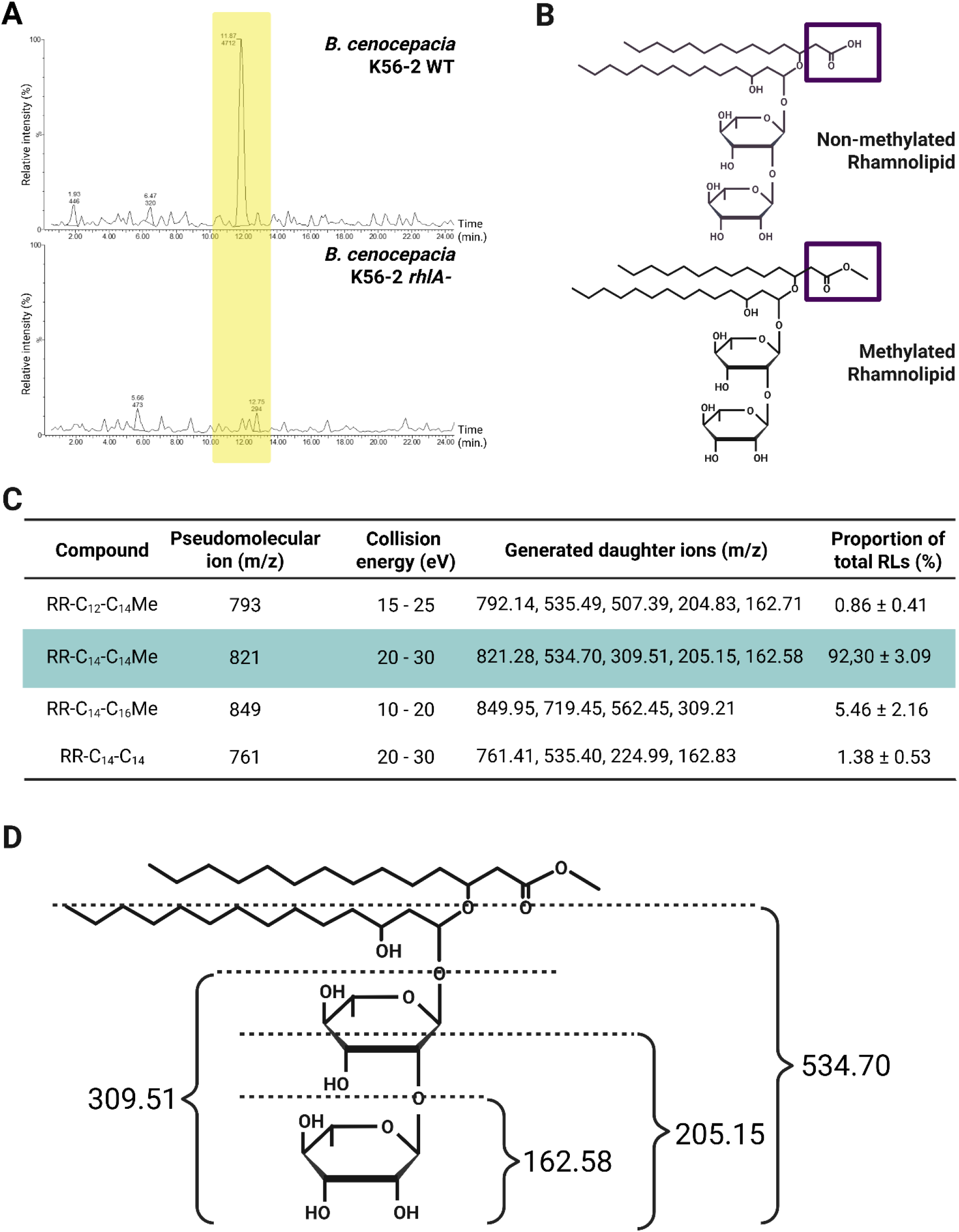
Identification of methylated rhamnolipids produced by *Burkholderia cenocepacia* K56-2. (A) Extracted ion chromatograms at *m/z* 821.6 for *B. cenocepacia* K56-2 wild-type (WT) versus the *rhlA* mutant. (B) Chemical structures of a standard non-methylated di-rhamnolipid (top) and a methylated di-rhamnolipid congener (bottom). The violet boxes highlight the structural difference at the carboxyl terminus (free carboxylic acid vs. methyl ester group). (C) Table summarizing the pseudomolecular ions (*m*/*z*) and corresponding generated daughter ions (*m*/*z*) obtained by LC-MS/MS analysis for the different methylated (RR-CxCyme) and non-methylated (RR-CxCy) rhamnolipid congeners detected. The proportion of each congener relative to total rhamnolipids was estimated from its LC-MS peak area and expressed as a percentage of the summed peak areas of all detected rhamnolipid congeners. The predominant congener is highlighted in blue. (D) Proposed MS/MS fragmentation pattern of a representative methylated dirhamnolipid congener. Dashed lines indicate the predicted cleavage sites during collision-induced dissociation corresponding to the observed daughter ions in (C).

To further confirm their identity, tandem mass spectrometry (LC-MS/MS) was performed. The product ion analysis (daughter ions) of the fragmented molecules revealed specific cleavage pathways (**Fig. 2C and 2D**), confirming that these compounds are indeed different congeners of methylated rhamnolipids. Unlike rhamnolipids, these molecules possess a methyl ester group at their carboxyl terminus (**Fig. 2B**), matching the unique structural profile we previously found in *B. lata* cultures (23). Interestingly, while the methylated forms were systematically detected as formate adducts [M + FA – H]^-^, the non-methylated congener did not form a formate adduct. This distinct ionization behavior complicated their initial detection and may also partly explain why rhamnolipid production in *B. cenocepacia* K56-2 had remained elusive. This quantification method allows us to further optimize production conditions for future applications.

Taken together, our data reveal that the biosurfactant produced by *B. cenocepacia* K56-2 is composed of rhamnolipids and successfully determine that the main congeners are methylated, and thus non-ionic. In all evidence, this explains why the biosurfactant produced by *B. cenocepacia* had remained elusive.

### Widespread distribution and functional conservation of the *rhlM*-containing operon across the Burkholderia cepacia complex

Many species of the *Burkholderiaceae* family possess the *rhl* operon, and most of them also carry the *rhlM* gene (13), suggesting that these potential rhamnolipid producers may also produce methylated rhamnolipids. Intriguingly, a large proportion of these potential producers belong to the Bcc (13), including *B. cenocepacia* (this study) and *B. lata* (23), two species now confirmed to produce methylated rhamnolipids. To further investigate the distribution of this biosynthetic potential across the Bcc, we examined the presence and organization of the *rhl* operon across the currently identified Bcc species. Of these, 19 had at least one sequenced genome available for analysis. The *rhl* operon was identified in 15 of these 19 species (78.95%), while it was absent from all available genomes of the remaining four species: *Burkholderia dolosa*, *Burkholderia multivorans*, *Burkholderia ubonensis* and *Burkholderia vietnamiensis*.For most *rhl*-positive species, the operon was detected in all available genomes, although some variation was observed, notably in *Burkholderia ambifaria* (9/10 genomes) and *Burkholderia contaminans* (18/22 genomes). Importantly, in all genomes in which the operon is present, the *rhlM* gene is always found between *rhlA* and *rhlB* in the biosynthetic gene cluster (13, 22) (**Table 5**).

**Table 5.**
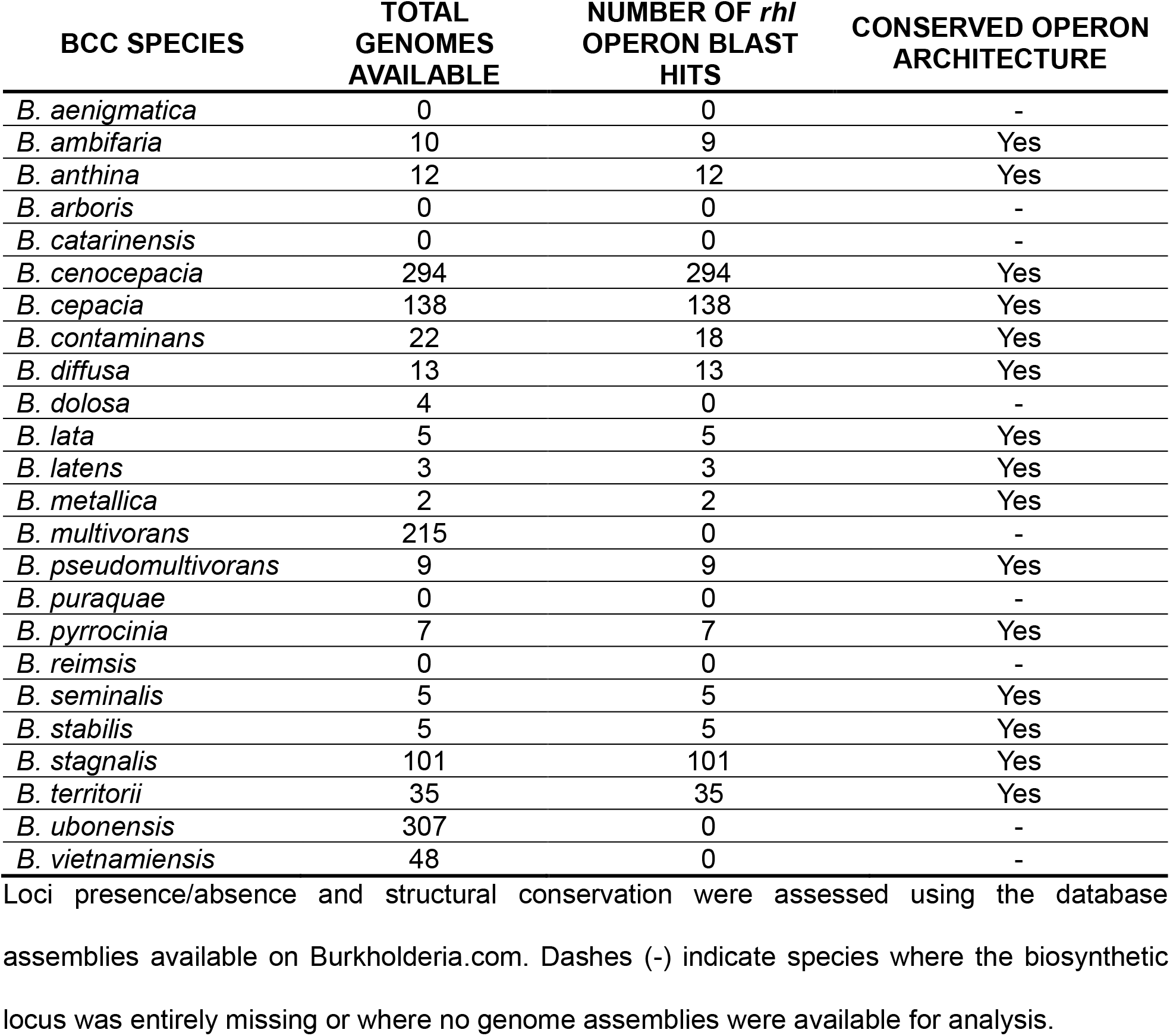
Distribution and genomic conservation of the *rhl* operon across *Burkholderia cepacia* complex species.

| BCC SPECIES | TOTAL GENOMES AVAILABLE | NUMBER OF *rhI* OPERON BLAST HITS | CONSERVED OPERON ARCHITECTURE |
| --- | --- | --- | --- |
| *B. aenigmatica* | 0 | 0 | - |
| *B. ambifaria* | 10 | 9 | Yes |
| *B. anthina* | 12 | 12 | Yes |
| *B. arboris* | 0 | 0 | - |
| *B. catarinensis* | 0 | 0 | - |
| *B. cenocepacia* | 294 | 294 | Yes |
| *B. cepacia* | 138 | 138 | Yes |
| *B. contaminans* | 22 | 18 | Yes |
| *B. diffusa* | 13 | 13 | Yes |
| *B. dolosa* | 4 | 0 | - |
| *B. lata* | 5 | 5 | Yes |
| *B. latens* | 3 | 3 | Yes |
| *B. metallica* | 2 | 2 | Yes |
| *B. multivorans* | 215 | 0 | - |
| *B. pseudomultivorans* | 9 | 9 | Yes |
| *B. puraquae* | 0 | 0 | - |
| *B. pyrrocinia* | 7 | 7 | Yes |
| *B. reimsis* | 0 | 0 | - |
| *B. seminalis* | 5 | 5 | Yes |
| *B. stabilis* | 5 | 5 | Yes |
| *B. stagnalis* | 101 | 101 | Yes |
| *B. territorii* | 35 | 35 | Yes |
| *B. ubonensis* | 307 | 0 | - |
| *B. vietnamiensis* | 48 | 0 | - |

Loci presence/absence and structural conservation were assessed using the database assemblies available on Burkholderia.com. Dashes (-) indicate species where the biosynthetic locus was entirely missing or where no genome assemblies were available for analysis.

To determine whether the presence of the *rhl* operon, including the *rhlM* gene, correlates with the production of methylated rhamnolipids, cultures of at least one strain from 18 Bcc species, either carrying or lacking the *rhl* operon, were prepared using the optimized culture conditions we developed for *B, cenocepacia* K56-2. Although productivity varied among strains, methylated rhamnolipids were detected in all strains in which the operon was identified. In all cases, the dominant congener detected was RR-C_14_-C_14_Me. Several strains also produced the same additional congeners, such as RR-C_12_-C_14_Me and RR-C_14_-C_16_Me, that we identified in cultures of *B. cenocepacia* K56-2 (**Fig. 3A** and **Fig. 4**).

**Figure 3.**
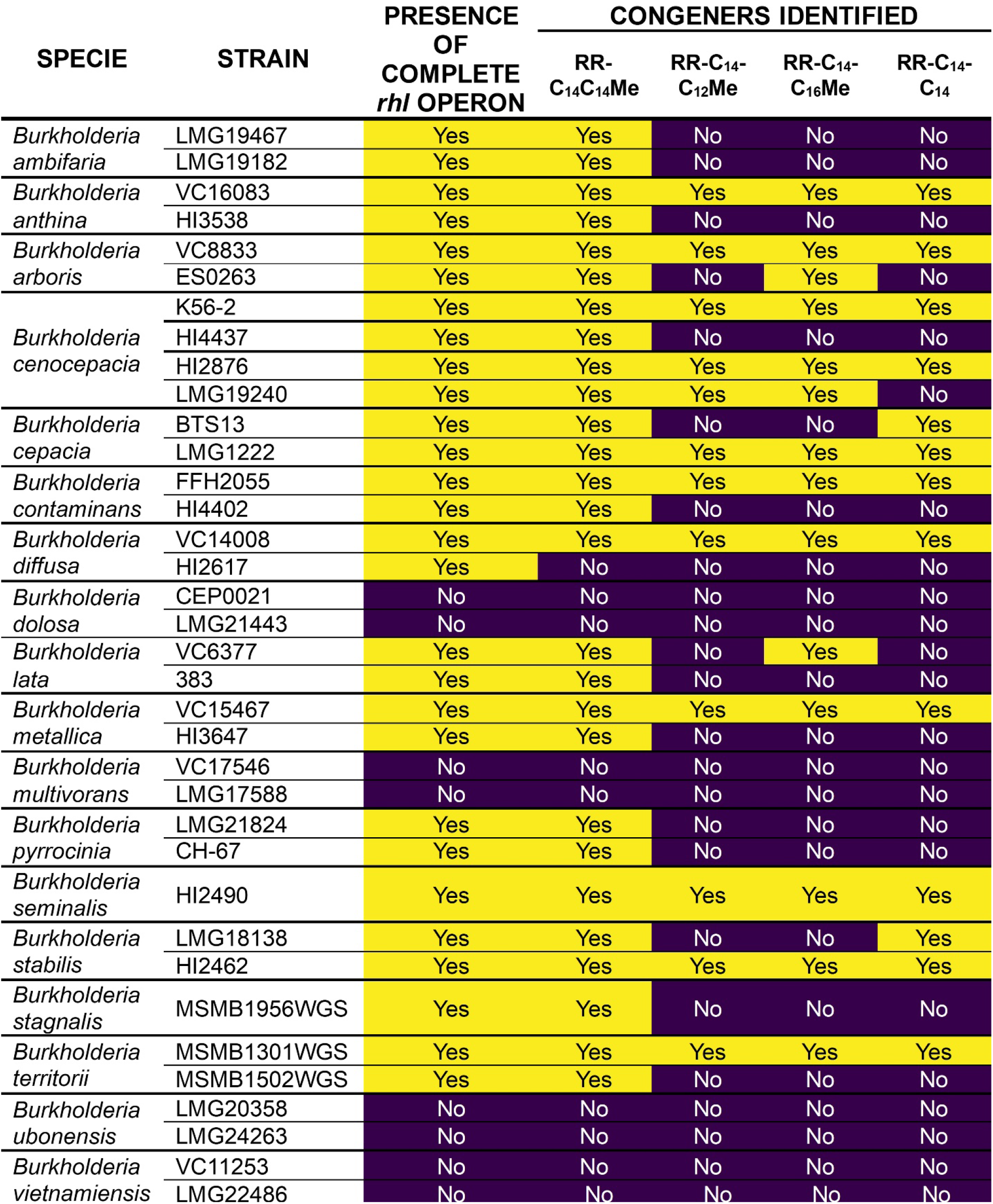
Distribution of identified methylated (RR-CXCYMe) and non-methylated (RR-CXCY) rhamnolipid congeners across Bcc strains. Presence of the complete *rhl* operon and detection of individual rhamnolipid congeners were assessed across the Bcc strains analyzed in this study. Yellow and purple cells indicate presence/detection and absence/non-detection, respectively. Congeners detected included RR-C_14_C_14_Me, RR-C_14_-C_12_Me, RR-C_14_-C_16_Me and RR-C_14_-C_14_.

**Figure 4.**
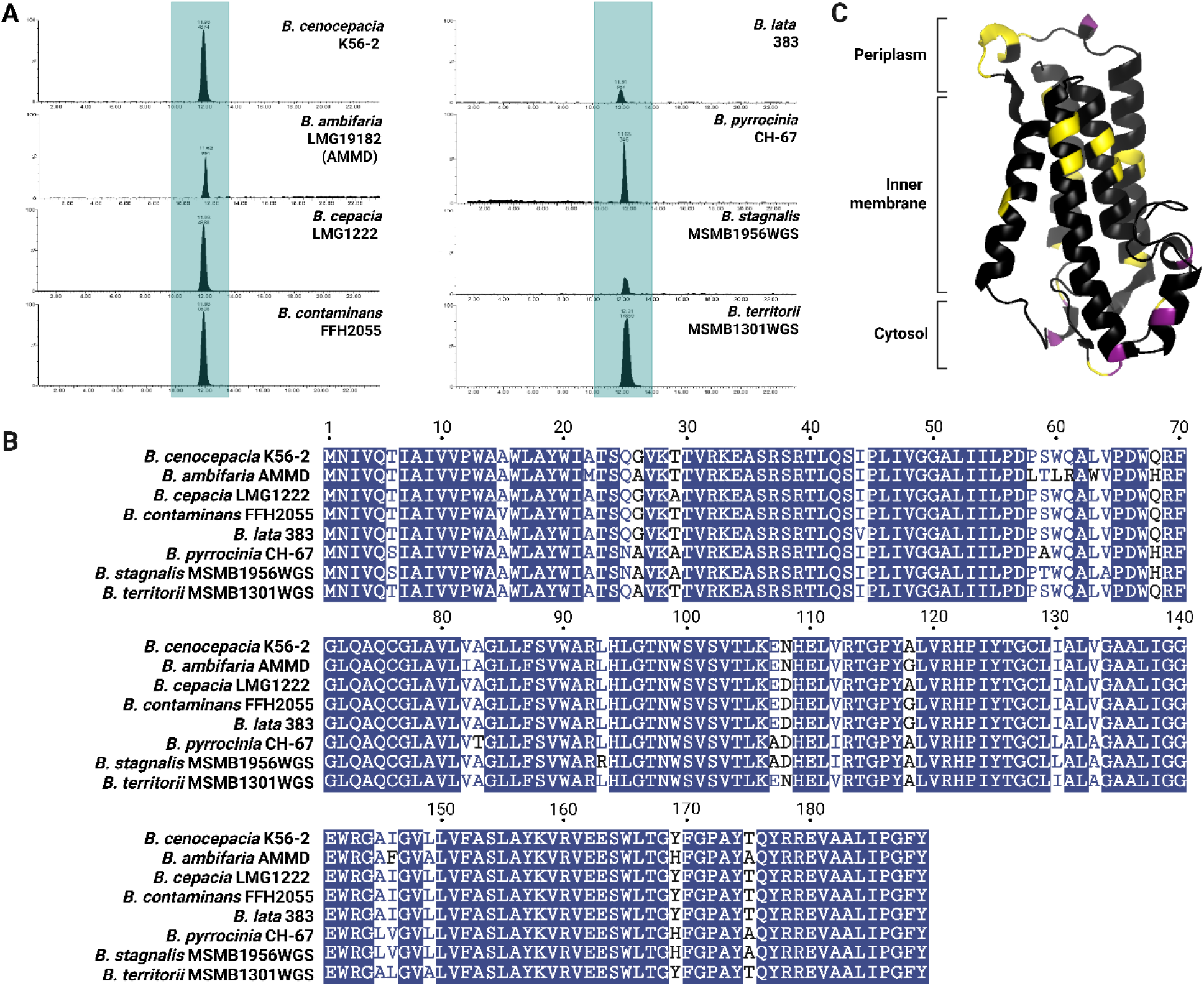
Genomic conservation and structural analysis of RhlM homologs across the Bcc. (A) Representative LC-MS extracted ion chromatograms showcasing the production of methylated rhamnolipids across diverse *rhl*-positive Bcc member species grown under optimized conditions. (B) Multiple amino acid sequence alignment of selected Bcc RhlM homologs, highlighting a high degree of conservation with limited amino acid substitutions. (C) Mapping of sequence variability onto the AlphaFold-predicted 3D structure of the *B. cenocepacia* K56-2 RhlM protein. Substitutions are colored to distinguish between conservative changes with minor predicted structural impacts (yellow) and more divergent substitutions (green) located near the predicted cytosolic catalytic site.

These results successfully confirm and clarify several historical suspicions regarding biosurfactant production within the Bcc (4–7). Notably, a previous study identified a C_15_-C_14_ dirhamnolipid congener in cultures of *Burkholderia seminalis* (3). Given that bacterial fatty acid synthesis predominantly yields even-numbered carbon chains, such an odd-numbered C_15_ fatty acid architecture is highly unusual (20, 60). Our findings strongly suggest that this previously reported molecule was likely a misidentified methylated C_14_-C_14_ congener, where the extra carbon originates from the RhlM-mediated methyl ester group rather than a typical longer lipid chain. Overall, this shows for the first time that rhamnolipid production is a widespread trait among Bcc species and that the major congeners produced are methylated, likely thanks to the presence of the recently identified methyltransferase RhlM.

### Genomic conservation and structural modeling of the RhlM methyltransferase

Since the standout feature of rhamnolipid-producing Bcc species is the methylation, we further investigated the *rhlM* gene. To further assess genomic conservation, a BLASTn alignment was performed for strains with available genome sequences, using the *rhlM* gene sequence from *B. cenocepacia* K56-2 as a reference. In all strains where *rhlM* was identified, homologues of *rhlA*, *rhlB*, and *rhlC* were also present and maintained the same genomic organization as in *B. cenocepacia* K56-2 (**Fig. 4B**).

Alignment of the predicted RhlM amino acid sequences revealed that the protein is highly conserved across the Bcc strains analyzed. The length of the protein is identical in all strains, and only a small number of amino acid substitutions were identified (**Fig. 4B**). When these substitutions were mapped onto the AlphaFold-predicted structure of RhlM from *B. cenocepacia* K56-2, the protein appeared largely conserved, with substitutions confined to a limited number of positions. Some substitutions correspond to conservative changes with minor predicted structural impact (yellow), whereas others correspond to more divergent substitutions (green) (**Fig. 4C**).

Interestingly, the transmembrane helices were almost entirely conserved among the strains, while the few substitutions predicted to have a larger impact were located in the cytosolic portion of the protein. As proposed by Liu *et al*. (22), the active site of the RhlM enzyme is predicted to be located on the cytosolic side of the membrane, as in other membrane-bound methyltransferases, such as the isoprenylcysteine carboxyl methyltransferases from *Methanosarcina acetivorans* and *Tribolium castaneum* (22). In this model, the cofactor pocket that binds S-adenosylmethionine is positioned on the cytosolic side, whereas a substrate located closer to the transmembrane region provides access to the lipid substrate (22).

Mapping the sequence variability observed among the Bcc RhlM homologs onto this structural model suggests that the variable residues are located near the catalytic region (**Fig. 4C**). These substitutions could therefore influence several aspects of enzyme function, including substrate recognition, substrate binding, accessibility of the lipid tunnel, or catalytic efficiency (95).

However, since several congeners of methylated dirhamnolipids were detected across all strains analyzed (**Fig. 3**, **Fig. 4A**), these variations do not appear to prevent the methylation of different rhamnolipid congeners. This suggests that RhlM can accommodate some structural variation in its substrates, at least among the long-chain rhamnolipids produced by Bcc species. These variations could potentially affect enzyme activity and contribute to the differences in methylated rhamnolipid production observed between strains. Moreover, this substrate flexibility demonstrates that RhlM possesses an adaptable catalytic pocket, capable of accommodating structural variations among long-chain lipids. Ultimately, this enzymatic tolerance establishes RhlM as a promising biocatalytic tool for engineering custom, non-ionic biosurfactants for industrial applications.

## CONCLUSION

In summary, our study demonstrates that rhamnolipid methylation is a structurally conserved and widespread trait across the Bcc, rather than an isolated structural exception. Indeed, rhamnolipids are extensively investigated biosurfactants, but are produced by only a handful of well-characterized bacterial species (20). In contrast, we found here that a large diversity of *Burkholderiaceae* apparently also possess the ability to produce rhamnolipids, but that these are methyl esters. By optimizing culture conditions using a bioluminescent reporter system, we successfully enhanced *rhl* operon expression in *B. cenocepacia* K56-2, identifying nutritional conditions triggering higher expression. These findings open fundamental questions regarding the evolutionary path of rhamnolipid producers. Indeed, genomic data indicate that the *rhlM* gene is more often present than absent among producing *Burkholderiaceae* (13), suggesting that methylated rhamnolipids may represent the ancestral form of these biosurfactants. Consequently, non-methylating producers, such as *P. aeruginosa* or *B. thailandensis*, might have lost the *rhlM* gene during their evolutionary diversification. However, this hypothesis should be interpreted cautiously, as many *rhlM*-containing species belong to the closely related Bcc (13, 22), which may contribute disproportionately to the apparent prevalence of the gene.

This evolutionary divergence raises a pivotal question: Why did certain bacteria evolve to maintain rhamnolipid methylation while others discarded it? As proposed by Liu *et al*. (22) , the functional purpose of methylating rhamnolipids may be linked to specific environmental adaptation, as neutralizing the negative charge of the carboxyl terminus to produce non-ionic surfactants could provide distinct ecological advantages. Characterizing how this structural modification alters the physicochemical and biological behaviors of these molecules will be a crucial next step to understand their biological significance. This question is particularly relevant in the Bcc, whose members are opportunistic pathogens (1). Biosurfactants to contribute to traits involved in environmental colonization and persistence, as well as interactions with hosts and virulence (12). Rhamnolipid methylation could therefore modulate these functions, although its specific contribution remains unknown. Intriguingly, the absence of the *rhl* operon in some Bcc species, including *B. dolosa*, *B. multivorans*, *B. ubonensis*, and *B. vietnamiensis*, provides a natural contrast that may help determine the ecological and biological advantages conferred by rhamnolipid production and methylation. Understanding these functions will also be important for evaluating the biotechnological potential of methylated rhamnolipids. Ultimately, shifting the paradigm of rhamnolipid diversity from an exception to the rule underscores the necessity to unveil both the functional properties and the ecological drivers behind this widespread structural modification.

## ACKNOWLEDGEMENTS

We thank Danielle Dubeau for initiating this long-standing project and Donat Monoko for the construction of the *rhlA* mutant. We also acknowledge Margaux Lustig for her contribution to this work. We are grateful to Sylvain Milot and Marianne Piochon for their valuable technical assistance with biochemical extractions and LC-MS analyses throughout this project. Finally, we sincerely thank Dr. Sonja Kubicki for her insightful discussions and collaboration in the structural identification of methylated rhamnolipids.

