## Supplemental material for "From Exception to Rule: Widespread Production of Methylated Rhamnolipids in the *Burkholderia cepacia* Complex"

### SUPPLEMENTARY METHODS

#### ***In silico* screening of the *rhl* operon across the Bcc**

To evaluate the distribution and conservation of the *rhl* biosynthetic locus within *the Burkholderia cepacia* complex (Bcc), an *in silico* screening was performed using the *Burkholderia* Genome Database (burkholderia.com). The nucleotide sequence of the *rhlM* gene from *B. cenocepacia* K56-2 was used as a query for high-stringency BLASTn searches against all available reference and draft assemblies for each described Bcc species. To avoid false positives resulting from non-specific short alignments or homologies with other methyltransferases, only full-length hits covering the complete marker gene (~570 pb) with high identity were counted as positive. The genetic organization of the flanking regions was manually inspected to confirm the conservation of the *rhlA-rhlM-rhlB* gene synteny.

### SUPPLEMENTARY RESULTS

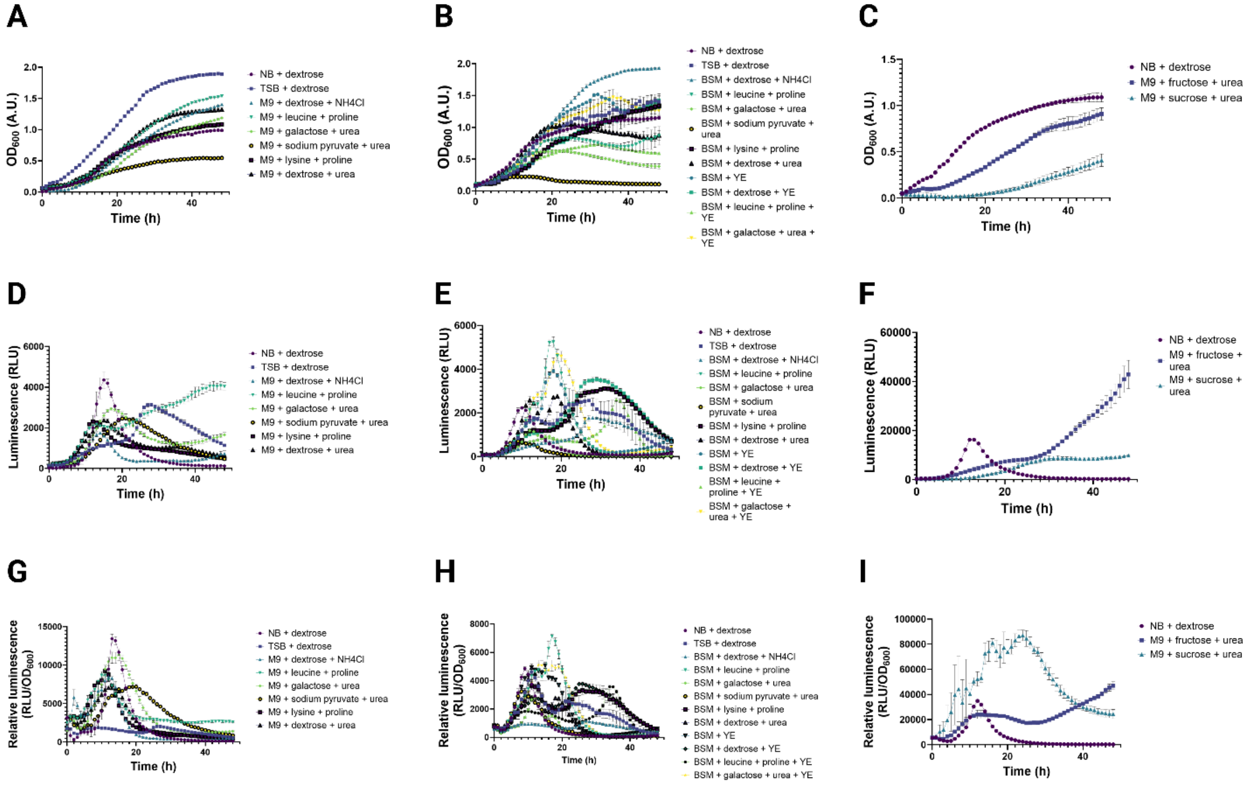


**Figure S1. Growth and P*rhlI* reporter activity profiles of *B. cenocepacia* K56-2 under the culture conditions evaluated during the primary screening.** Growth ($OD_{600}$), bioluminescence (RLU), and relative P*rhlI* activity ($RLU/OD_{600}$) were monitored over 48 hours in 96-well plates. (A-C) Growth profiles, (D-F) corresponding bioluminescence profiles, and (G-I) relative P*rhlI* activity profiles. NB supplemented with dextrose was included as a reference condition. Data are presented as mean ± SD of four biological replicates.

**Table S1. Growth and *rhl* operon expression parameters obtained during the initial culture medium screening.**

| Medium ID | $AUC_{OD600}$ (A.U. • h) | $AUC_{lum}$ (RLU • h) | $AUC_{rel.lum}$ (RLU/A.U. • h) | $AUC_{combined.inedx}$ (RLU x A.U. • h) |
|---|---|---|---|---|
| **1** | 28.70 ± 1.15 | 43604 ± 3711 | 115442 ± 4929 | 21933 ± 2522 |
| **2** | 56.54 ± 0.29 | 75039 ± 1596 | 67743 ± 886 | 110085 ± 2883 |
| **3** | 32.89 ± 0.29 | 26792 ± 586 | 108405 ± 4266 | 16686 ± 723 |
| **4** | 38.19 ± 0.15 | 115317 ± 4497 | 183011 ± 4618 | 119298 ± 5853 |
| **5** | 28.23 ± 0.37 | 69113 ± 1829 | 186188 ± 6781 | 42821 ± 1838 |
| **6** | 17.04 ± 0.37 | 57058 ± 1329 | 173832 ± 3340 | 22110 ± 739 |
| **7** | 30.87 ± 0.60 | 47207 ± 538 | 126690 ± 1945 | 28447 ± 648 |
| **8** | 36.55 ± 1.03 | 53008 ± 1084 | 119919 ± 5549 | 38988 ± 1038 |
| **9** | 17.92 ± 0.94 | 178032 ± 13395 | 424029 ± 10464 | 108413 ± 15468 |
| **10** | 5.11 ± 0.71 | 71357 ± 6049 | 912164 ± 52647 | 12675 ± 3015 |
| **11** | 40.85 ± 0.32 | 86677 ± 1453 | 102761 ± 2493 | 105459 ± 1407 |
| **12** | 21.27 ± 1.20 | 107297 ± 2836 | 249680 ± 4652 | 58283 ± 2524 |
| **13** | 15.55 ± 0.66 | 24848 ± 2032 | 78259 ± 3900 | 10019 ± 1126 |
| **14** | 5.33 | 15557 | 109953 | 2535 |

| | | | | |
|---|---|---|---|---|
| | ± 0.19 | ± 51 | ± 4885 | ± 115 |
| **15** | 26.00 ± 2.20 | 156200 ± 2785 | 290030 ± 19824 | 115492 ± 10207 |
| **16** | 26.20 ± 0.48 | 76703 ± 997 | 163468 ± 2787 | 48639 ± 601 |
| **17** | 32.01 ± 0.21 | 100503 ± 2273 | 226098 ± 5448 | 72269 ± 1914 |
| **18** | 26.54 ± 0.40 | 170170 ± 6411 | 299713 ± 14096 | 128821 ± 3154 |
| **19** | 18.11 ± 0.12 | 78179 ± 1582 | 194274 ± 2783 | 38923 ± 1034 |
| **20** | 31.88 ± 1.21 | 125804 ± 2823 | 247794 ± 1207 | 87673 ± 5314 |

Medium IDs correspond to those presented in Figure 1A. Values are presented as mean ± standard deviation of 4 replicates. $AUC_{OD600}$, area under the $OD_{600}$ curve; $AUC_{lum}$, area under the luminescence (RLU) curve; $AUC_{rel.lum}$, area under the relative luminescence (RLU/$OD_{600}$) curve; $AUC_{combined.index}$, combined index calculated from $OD_{600}$ and luminescence AUCs. $OD_{600}$, optical density at 600 nm; RLU, relative light units; A.U., arbitrary units.

**Table S2. Growth and *rhl* operon expression parameters obtained during the characterization of selected culture conditions.**

| | 1 | 2 | 3 | 4 | 5 | 6 | 7 | 8 |
|---|---|---|---|---|---|---|---|---|
| $\mu_{max}$ (h-1) | 0.222 ± 0.004 | 0.787 ± 0.087 | 0.316 ± 0.010 | 0.363 ± 0.044 | 0.288 ± 0.010 | 0.712 ± 0.058 | 0.196 ± 0.016 | 0.191 ± 0.017 |
| G (h) | 3.124 ± 0.060 | 0.888 ± 0.102 | 2.194 ± 0.071 | 1.925 ± 0.221 | 2.405 ± 0.082 | 0.978 ± 0.077 | 3.560 ± 0.302 | 3.650 ± 0.322 |
| $P_{OD600.max}$ (A.U./L/h) | 0.352 ± 0.044 | 0.999 ± 0.441 | 0.271 ± 0.001 | 0.755 ± 0.066 | 0.308 ± 0.010 | 0.331 ± 0.018 | 0.328 ± 0.009 | 0.389 ± 0.012 |
| $P_{OD600.tot}$ (A.U./L/h) | 0.320 ± 0.16 | 0.358 ± 0.09 | 0.271 ± 0.001 | 0.647 ± 0.034 | 0.308 ± 0.010 | 0.322 ± 0.018 | 0.308 ± 0.010 | 0.380 ± 0.023 |
| $P_{lum.max}$ (RLU/L/h) | 1016 ± 43 | 341 ± 24 | 2881 ± 264 | 3049 ± 541 | 1853 ± 296 | 3350 ± 385 | 4665 ± 279 | 1669 ± 107 |
| $P_{lum.tot}$ (RLU/L/h) | 256 ± 12 | 39 ± 4 | 2881 ± 264 | 1220 ± 220 | 1640 ± 82 | 783 ± 18 | 3045 ± 35 | 1669 ± 107 |
| $P_{lum.rel.max}$ (RLU/A.U./L/h) | 264 ± 63 | 54 ± 4 | 4651 ± 455 | 2222 ± 1183 | 3872 ± 335 | 727 ± 139 | 1490 ± 90 | 1625 ± 369 |
| $P_{lum.rel.tot}$ (RLU/A.U./L/h) | 47 ± 4 | 6 ± 1 | 626 ± 57 | 111 ± 21 | 313 ± 6 | 144 ± 11 | 584 ± 20 | 259 ± 9 |
| $AUC_{DO600}$ (A.U. • h) | 50.62 ± 0.78 | 91.06 ± 1.87 | 19.31 ± 0.49 | 80.84 ± 4.54 | 22.48 ± 0.25 | 63.60 ± 1.60 | 38.79 ± 0.90 | 38.47 ± 0.92 |
| $AUC_{RLU}$ (RLU • h) | 88505 ± 1729 | 22014 ± 430 | 348871 ± 10541 | 259117 ± 15781 | 253397 ± 2855 | 282225 ± 3216 | 424301 ± 6266 | 246293 ± 7742 |
| $AUC_{rel.\ lum}$ (RLU/U.A. • h) | 35281 ± 5223 | 10907 ± 3886 | 333424 ± 15426 | 108013 ± 15254 | 274416 ± 13023 | 77165 ± 3435 | 151033 ± 3990 | 134379 ± 6856 |

Eight culture conditions were evaluated: (1) NB + dextrose, (2) TSB + dextrose, (3) M9 + galactose + urea, (4) M9 + fructose + urea, (5) M9 + leucine + proline, (6) BSM + dextrose + urea, (7) BSM + galactose + urea, and (8) BSM + lysine + proline. Values are presented as mean ± standard deviation of 3 replicates. $\mu_{max}$, maximum specific growth rate; G, generation time; $P_{OD600,max}$, maximum $OD_{600}$ productivity; $P_{OD600,tot}$, total $OD_{600}$ productivity; $P_{lum,max}$, maximum luminescence productivity; $P_{lum,tot}$, total luminescence productivity; $P_{lum,rel,max}$, maximum relative luminescence productivity; $P_{lum,rel,tot}$, total relative luminescence productivity; $AUC_{OD600}$, area under the $OD_{600}$ curve; $AUC_{RLU}$, area under the luminescence curve; $AUC_{rel.lum}$, area under the relative luminescence (RLU/$OD_{600}$) curve. $OD_{600}$, optical density at 600 nm; RLU, relative light units; A.U., arbitrary units.

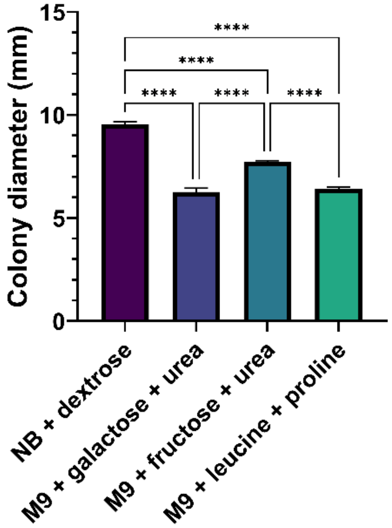


**Figure S2. Colony diameters of *Burkholderia cenocepacia* K56-2 across selected culture media.** Bacteria were grown on agar plates prepared with the four indicated media for 48 hours at 30°C. Data are presented as mean ± SD of three biological replicates. Statistical significance was determined using a one-way ANOVA followed by Tukey's post-hoc test for multiple comparisons ($p < 0.0001$ = ****).

A

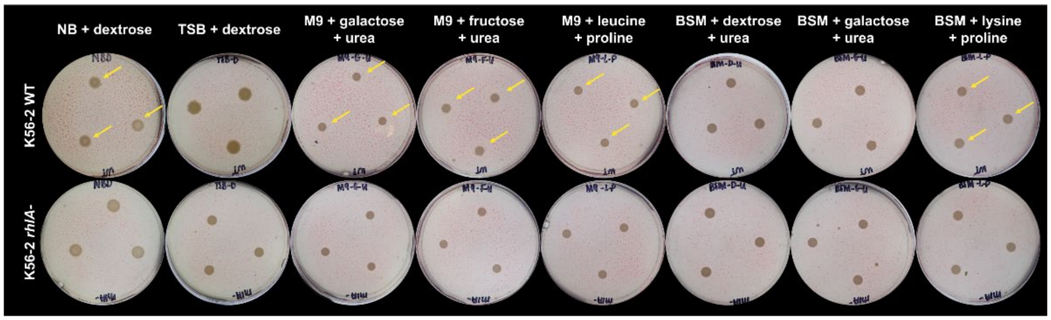


B

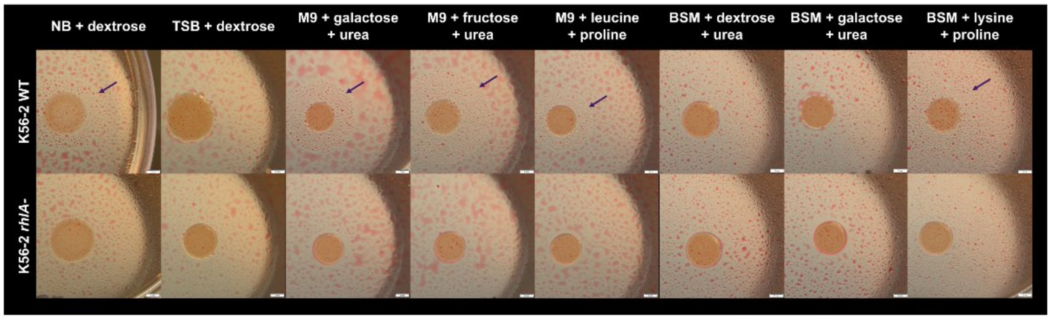


**Figure S3. Sprayed oil assay of *B. cenocepacia* K56-2 grown on selected culture media.** (A) Representative images of *B. cenocepacia* K56-2 wild type (WT) and the *rhlA*-deficient mutant grown on agar plates prepared with the eight culture media selected during the screening of *rhl* operon expression. Following incubation, plates were sprayed with mineral oil containing 0,5% Sudan Red to visualize oil displacement associated with biosurfactant production. Yellow arrows indicate representative oil displacement halos surrounding WT colonies. The *rhlA*-deficient mutant was included as a negative control for rhamnolipid production. (B) Representative microscopic images of the oil displacement zones surrounding WT and *rhlA*-deficient colonies under each culture condition. Purple arrows indicate the boundary of representative oil displacement halos.

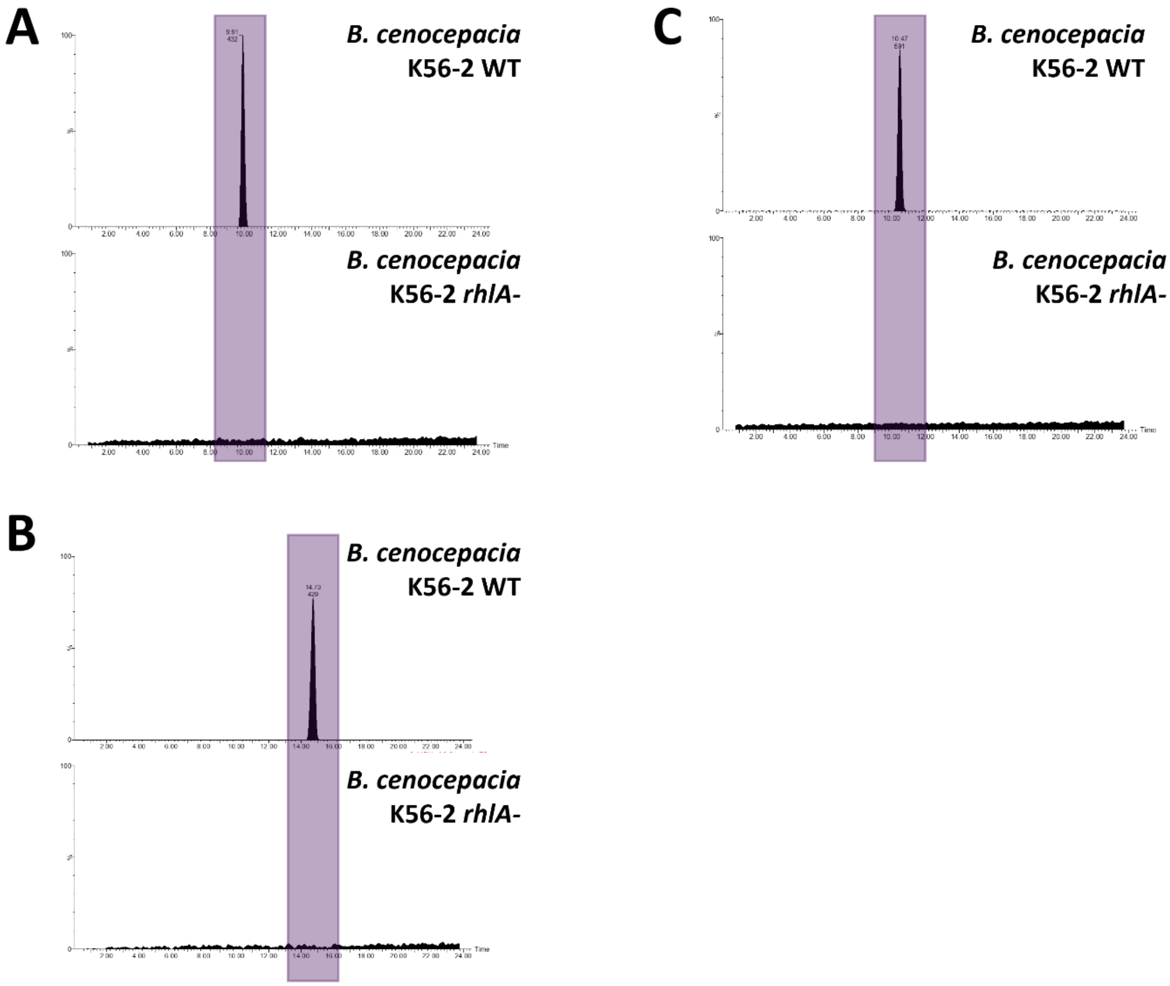


**Figure S4. Extracted ion chromatograms of the minor rhamnolipid congeners produced by *Burkholderia cenocepacia* K56-2.** Comparison between *B. cenocepacia* K56-2 WT, and the *rhlA* mutant cultures showing (A) the methylated congener at *m*/*z* 793 (meRR-$C_{12}$-$C_{14}$), (B) the methylated congener at *m*/*z* 849 (meRR-$C_{14}$-$C_{16}$), and (C) the non-methylated congener at *m*/*z* 761 (RR-$C_{14}$-$C_{14}$). All monitored peaks were strictly dependent on the rhamnolipid biosynthetic machinery, as evidenced by their complete absence in the *rhlA*-deficient mutant.

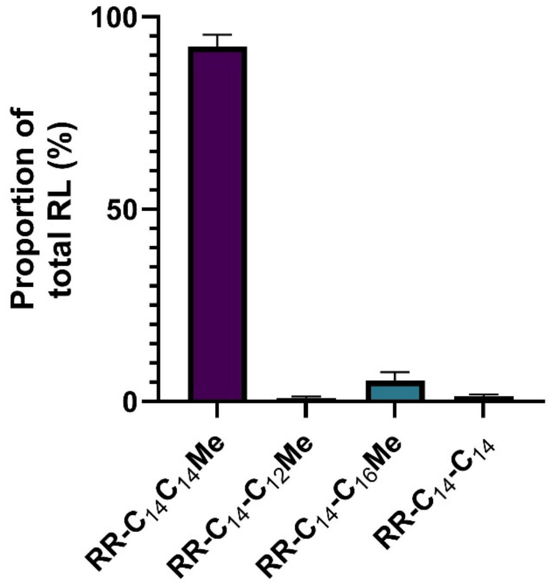


**Figure S5. Relative proportion of rhamnolipid congeners produced by *Burkholderia cenocepacia* K56-2.** For each biological replicate, the proportion of each congener was calculated from its LC-MS peak area relative to the summed peak areas of all detected rhamnolipid congeners. Data are presented as mean ± standard deviation.
